# Ultra-rare disruptive and damaging mutations influence educational attainment in the general population

**DOI:** 10.1101/050195

**Authors:** Andrea Ganna, Giulio Genovese, Daniel P. Howrigan, Andrea Byrnes, Mitja Kurki, Seyedeh M. Zekavat, Christopher W. Whelan, Mart Kals, Michel G. Nivard, Alex Bloemendal, Jonathan M. Bloom, Jacqueline I. Goldstein, Timothy Poterba, Cotton Seed, Robert E. Handsaker, Pradeep Natarajan, Reedik Mägi, Diane Gage, Elise B. Robinson, Andres Metspalu, Veikko Salomaa, Jaana Suvisaari, Shaun M. Purcell, Pamela Sklar, Sekar Kathiresan, Mark J. Daly, Steven A. McCarroll, Patrick F. Sullivan, Aarno Palotie, Tõnu Esko, Christina Hultman, Benjamin M. Neale

## Abstract

Ultra-rare inherited and *de novo* disruptive variants in highly constrained (HC) genes are enriched in neurodevelopmental disorders ^1–5^. However, their impact on cognition in the general population has not been explored. We hypothesize that disruptive and damaging ultra-rare variants (URVs) in HC genes not only confer risk to neurodevelopmental disorders, but also influence general cognitive abilities measured indirectly by years of education (YOE). We tested this hypothesis in 14,133 individuals with whole exome or genome sequencing data. The presence of one or more URVs was associated with a decrease in YOE (3.1 months less for each additional mutation; P-value=3.3×10^−8^) and the effect was stronger in HC genes enriched for brain expression (6.5 months less, P-value=3.4×10^−5^). The effect of these variants was more pronounced than the estimated effects of runs of homozygosity and pathogenic copy number variation ^6–9^. Our findings suggest that effects of URVs in HC genes are not confined to severe neurodevelopmental disorder, but influence the cognitive spectrum in the general population

---

Educational attainment, measured by the highest number of YOE attained is a complex trait influenced by public policy ^10^, economic resources^11^ and many heritable traits, including cognitive abilities and behavior ^12^. Importantly YOE is positively associated with healthy behaviors and lower rates of chronic diseases ^13–15^.

GWAS meta-analyses have identified 74 genome-wide significant loci for YOE ^16^. The additive heritability of YOE explained by common genetics variants has been estimated at 21% (95% C.I. 11-31%) ^17^,which is approximately half of the total heritability estimated from twin studies (40% 95% C.I. 35-44%) ^18^. It has been hypothesized that rare to ultra-rare exonic variants might account for some of the heritability currently not captured by GWAS ^19^.

Recent studies of intellectual disability, autism and schizophrenia have shed light on the impact of *de novo* and URVs on the genetic architecture of these disorders ^1–5^ (Genovese et al, *submitted*), showing a specific enrichment in HC genes (i.e. genes intolerant to loss-of-function or missense mutations). Moreover, emerging evidence suggests that *de novo* loss-of-function mutations are associated with reduced adaptive functioning in individuals without diagnosis of autism ^20^. Finally, using array data, a study has suggested that individuals with extremely high IQ have a reduced burden of rare protein-altering variants compared to unselected population-based controls^21^.

We tested the hypothesis that a burden of URVs in HC genes is associated with YOE in 14,133 individuals participating in four studies from three Northern European countries: Sweden, Estonia and Finland. Of these, 5,047 individuals have been diagnosed with schizophrenia. The remaining 9,086 individuals are selected to be free of schizophrenia or bipolar disorder, or from epidemiological studies, or from a biobank collection and are representative samples of the population (**Supplementary Material**). For each study we used the 1997 International Standard Classification of Education of the United Nations Educational, Scientific and Cultural Organization to define YOE.

The average numbers of YOE were 13.1, 13.6, and 11.8 in Swedish, Estonian, and Finnish participants, respectively. These differences are partially explained by different age and sex distributions, as well as by different methods used to measure educational attainment. For the Estonian and Finnish samples, we used self-report data; whereas for the Swedish sample’ we obtained YOE from the national registries (**Supplementary Table 1**).

We observed lower YOE among men (12.8 *vs.* 13.2 years, P-value=4.8×10^−12^) and older individuals (0.8 month less of education for each additional year of age, P-value<1×10^−15^) (**Supplementary Table 2**).

We developed a new software package called *Hail* to very efficiently perform quality control, annotation and analysis of large-scale sequencing data (**online Methods**).Using whole exome sequencing (WES) data (N. individuals=11,431) and protein coding regions in high-coverage whole genome sequencing (WGS) data (N. individuals=2,702), we identified URVs in HC genes (see the **Online Methods** for a detailed definition). URVs are variants that are observed only once (singletons) across each study and not observed in 60,706 exomes sequenced in the Exome Aggregation Consortium (ExAC) ^22^. The primary goal of this approach is to maximize the expected deleteriousness of the variants included (due to purifying selection). Within URVs we defined variants that were: (1) disruptive, putative loss-of-function variants including premature stop codons, essential splice site mutations and frameshift indels; (2) damaging, missense variants classified as damaging by seven different *in silico* prediction algorithms and (3) negative control, synonymous variants not predicted to change the encoded protein. We observed one or more of such mutations in 25%, 24%, 78% of individuals, respectively (**Supplementary Table 3**).

For each study, we fit a generalized linear regression model controlling for year of birth, sex, first 10 ancestry principal components, and schizophrenia status (**Online Methods**) to test for association of YOE with the number of disruptive or damaging URVs in HC genes (**Figure 1**) and meta-analyzed the results across studies. Principal components of genetic data showed that individuals within each study were of similar ancestry (**Supplementary Figure 1**).

On average, we observed a 3.1 months reduction in YOE for each disruptive mutation (P-value=3.3×10^−8^), and similar effect for damaging mutations (2.9 months less YOE, P-value=1.3×10^−6^). Furthermore, each additional disruptive mutation on average reduced the chance of going to college by 14% (odds ratio=0.86, P-value=0.0017). These results were consistent when using a mixed linear model approach to correct for population stratification in the Finnish and Estonian samples with WGS data (2.4 months less YOE; P-value=0.014, N=2,702).

The negative association between URVs and YOE remained consistent when we examined the control cohort and schizophrenia case cohort separately (**Supplementary Figure 2**). Furthermore, the effect remained consistent when excluding individuals diagnosed with a neurodevelopmental disorder (i.e. schizophrenia, bipolar disorder, autism, mental retardation and Asperger’s syndrome), as identified via linkage with the Swedish national inpatient registry (**Supplementary Figure 3**). We did not observe any significant association when we restricted our analysis to synonymous variants in HC genes (P-value=0.62) or disruptive mutations in unconstrained genes (P-value=0.73).

We used gene-expression data to determine whether restricting to genes enriched for brain expression concentrated our URVs burden signal. Specifically, we used the Genotype-Tissue Expression consortium data ^23^ to identify the 20% top brain-expressed HC genes. The intersection between HC and brain-expressed genes (N. genes=683 and 313 for disruptive and damaging URVs, respectively) more than doubled the impact on YOE (6.5 less months of YOE per each additional disruptive variant; P-value=3.4×10^−5^; **Figure 2**). The association was not significant when considering non brain-enriched HC genes or all brain-enriched genes (P-value>0.05). We further examined a subset of genes for which basal gene-expression was at least two fold higher in the brain compared to other tissues and observed similar results (6.3 less months of YOE; P-value=1×l0^−4^; **Supplementary Figure 4**)

To place disruptive and damaging URVs into context, we also examined the impact of previously reported genetic influences on YOE, including a polygenic score from common variants ^16^, runs of homozygosity ^6^,^7^ and a burden of rare pathogenic copy number variants (CNVs) ^8^,^9^. We sought to establish if these different forms of genetic variation act independently on YOE. For this purpose we defined four scores: (1) a polygenic score including all the independent single nucleotide polymorphisms (SNPs) with P-value<1 (as this threshold has been shown to maximize variance explained in YOE) in a large GWAS consortia of YOE ^16^ (2) the summed runs of homozygosity (3) disruptive and damaging URVs in HC genes and (4) self-curated list of pathogenic CNVs from the literature (**Supplementary Table 4**). The polygenic score was only calculated in the Swedish samples (N=10,644), since the other three studies were included in the original GWAS of YOE.

To assess the relative contribution of each genetic variation class to YOE, we fit the four normalized scores in the same regression model. All four scores were independently associated with YOE (**Figure 3**). The polygenic score showed the strongest association in standard deviations from the mean, explaining the largest proportion of the variability in YOE (2.9% *vs* 0.4% for the ultra-rare variants, 0.2% for runs of homozygosity and 0.1% for pathogenic CNVs).

We further evaluated whether the association between the polygenic score and YOE changes in individuals with and without disruptive or damaging URVs or CNVs. We found that the polygenic score was more strongly associated with YOE in individuals without disruptive or damaging URVs or CNVs (8.2 vs. 6.2 more months of YOE for 1 standard deviation increase in the polygenic score; P-value for interaction=0.007, **Supplementary Figure 5**).

Apart from genome-wide burden, we sought to identify individual genes driving the observed association between disruptive URVs and YOE. Using a gene-based burden test^24^ implemented in SKAT ^25^, and using an exome-wide significance threshold of 1×10^−6^, we didn’t identify any statistically significantly associated gene (**Supplementary Figure 6**).

In this study we focused on YOE, a phenotype that is relatively easy to collect in large samples and which has a strong genetic correlation with intelligence and cognitive function ^17^,^26^. We integrated WGS, WES and array data on more than 14,000 individuals and described the impact of URVs disrupting HC genes on YOE. This class of variants have been previously associated with autism^3^ and schizophrenia^4^, but the impact on cognition in the general population has not been described before. Here, for the first time, we show that disruptive and damaging URVs in HC genes are likely to affect cognition among individuals not diagnosed with neurodevelopmental disorders. Similar to the analyses of schizophrenia (Genovese et al, *submitted*) and autism, the majority of the signal lies in genes highly expressed in brain. This observation does not exclude the existence of causal mutations outside this gene class, but suggests that strong acting mutations are heavily concentrated within these genes.

Furthermore, we show that disruptive and damaging URVs in HC genes, common variants associated with YOE, runs of homozygosity, and CNVs implicated in monogenic syndromes or neurodevelopmental disorders, all act on cognitive function or personality traits ultimately reflected in the educational attainment of our study participants. This effect was not simply additive. We identified a modest, but significant interaction between the polygenic score and the presence of URVs or CNVs. Whether this observation is driven by the interplay of partially overlapping pathways between common and rare variants or by genotype-phenotype heterogeneity (e.g. common and rare variants impacting different subsets of individuals) will be a matter of future investigation.

Although stronger effect sizes were observed for CNVs and disruptive and damaging URVs, the polygenic score from common variants still explains the largest proportion of the YOE variability. This is not surprising, given that common variants are expected to have the largest contribution to heritable variation in most complex traits ^27^,^28^.

The prioritization approaches used to select variants contributing to the score from common variants and the score from rare variants are different. The former uses estimates of the association with YOE and the proportion of variance explained by the score is likely to improve once the sample size used to originate these estimates increases. The latter uses *in-silico* prediction of the variants’ functional effect coupled with population genetics expectations built on the mutation rate. As with the common variant score, we expect that this URV score will continue to improve in predictive validity of YOE as the characterization of which genes and genomic regions are associated with YOE further clarifies. For example, larger exome aggregation efforts would enable the calculation of exon-specific constraint scores or increases in exome sequencing on cohorts with YOE would further resolve which genes are relevant.

Our study could not detect disruptive or damaging mutations in a given gene as being unequivocally associated with YOE; however, as sample sizes increase, specific genes will emerge. Nevertheless, our proof-of-concept work shows that a wide range of genetic variation from ultra-rare disruptive mutations to CNVs and common variants influence cognitive function in the population.

**Fig. 1.**
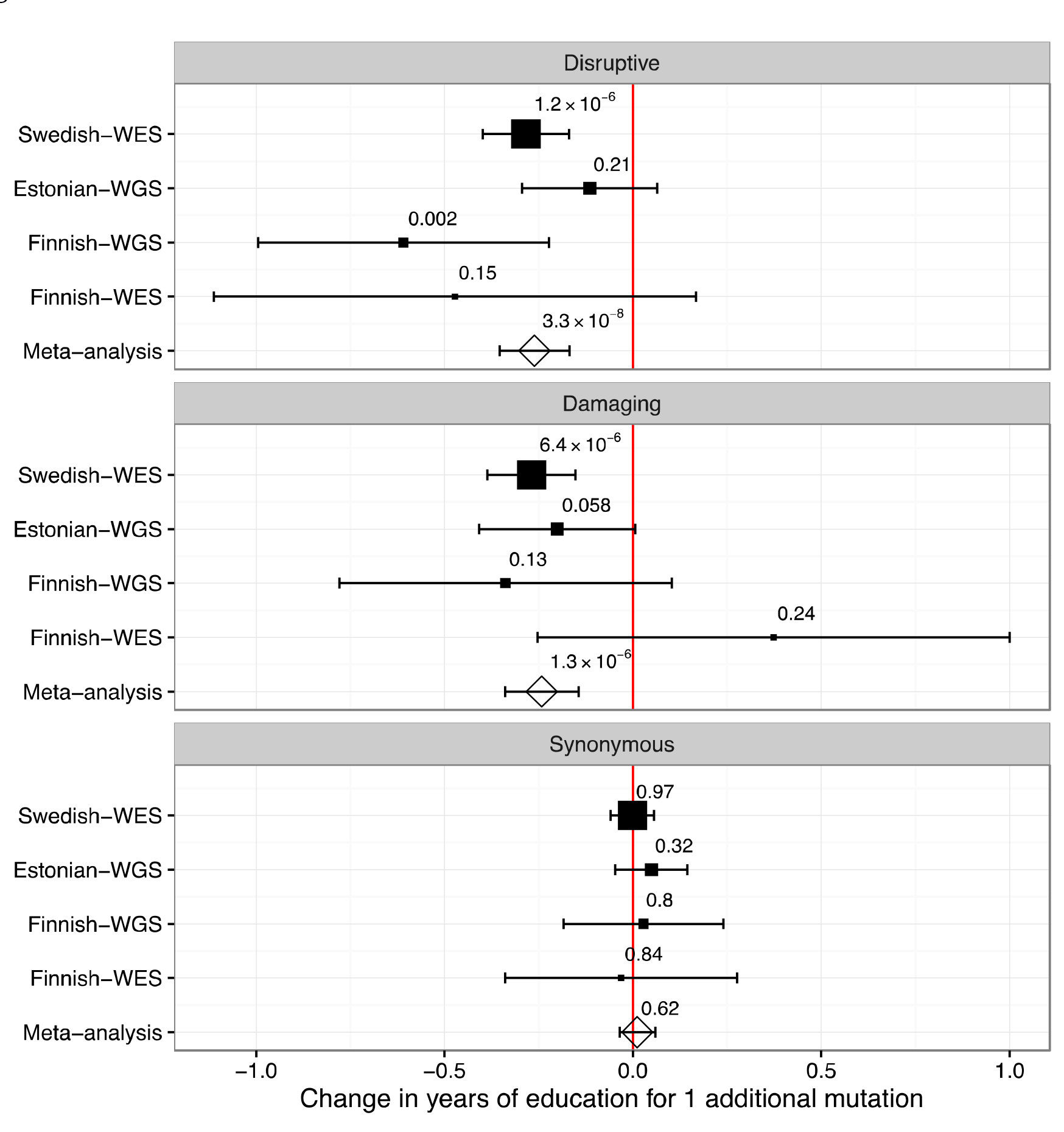
Forest plot for association between number of disruptive, damaging, and synonymous URVs in HC genes and YOE. The size of the squares is proportional to thesize of the study.

**Fig. 2.**
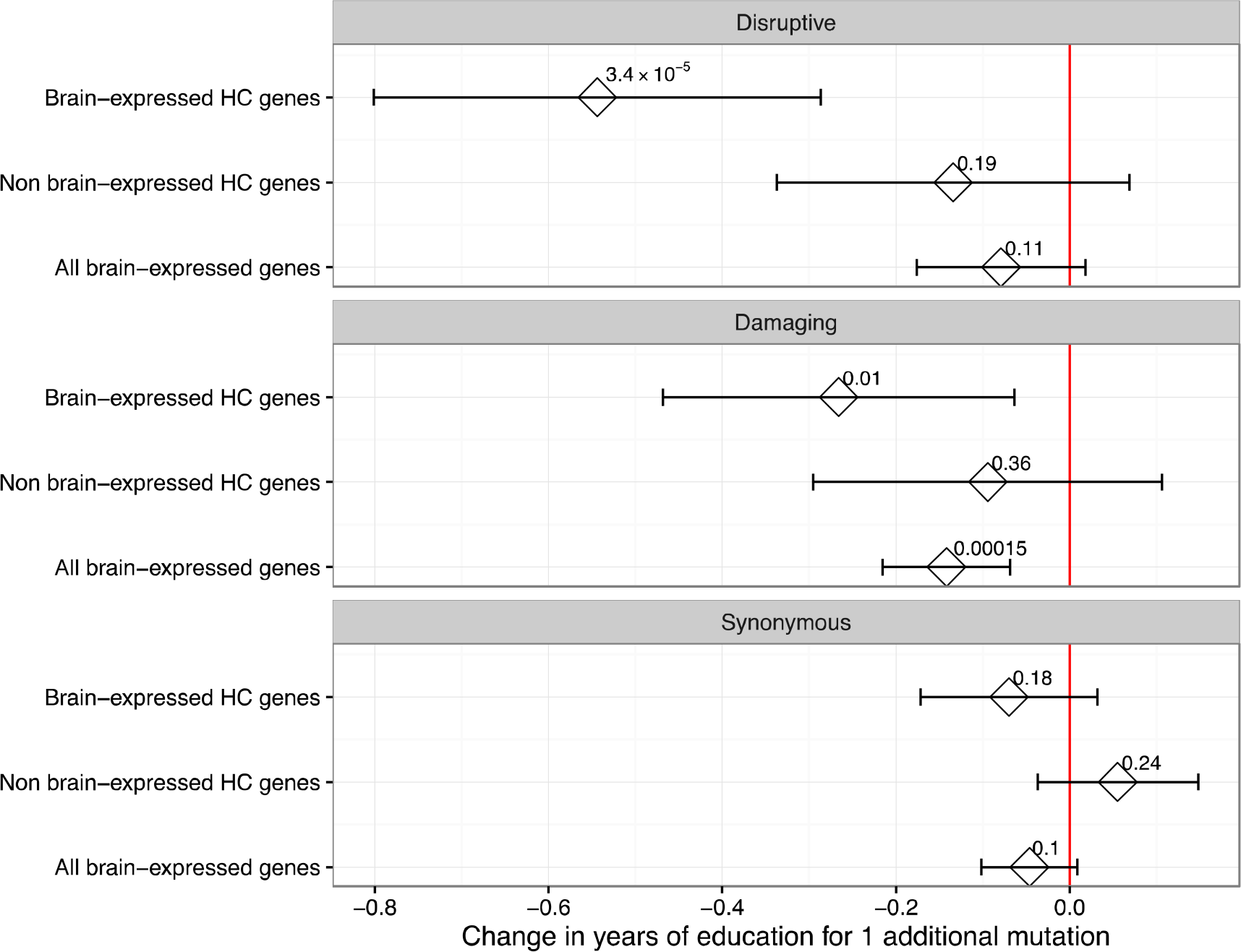
Association between numbers of disruptive, damaging and synonymous mutations for different gene sets. Meta-analysis results.

**Fig. 3.**
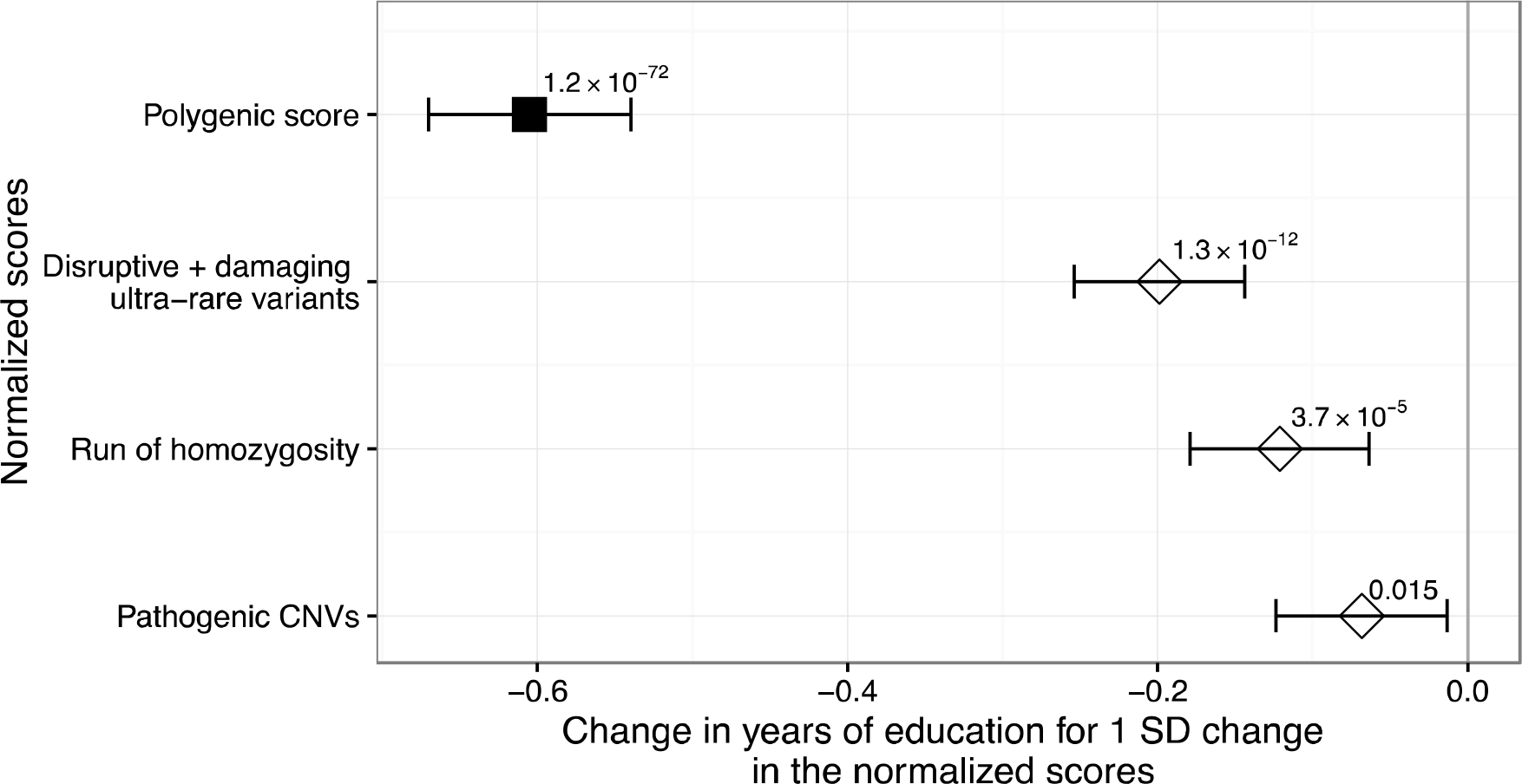
Association between each of the normalized scores (polygenic, runs of homozygosity, URVs and pathogenic CNVs) and YOE. The results presented are from meta-analysis of Swedish WES, Estonian WGS and Finnish WGS studies, except for the polygenic score, which is calculated only in the Swedish WES study. Notice that we plot 1-polygenic score to obtain a negative association with YOE.

URLs. Swedish WES data are available through dbGAP at http://www.ncbi.nlm.nih.gov/projects/gap/cgi-bin/study.cgi?studyid=phs000473

## ACKNOWLEDGMENT

This study was supported by grants from the National Human Genome Research Institute (U54 HG003067, R01 HG006855), the Stanley Center for Psychiatric Research, the Alexander and Margaret Stewart Trust, the National Institute of Mental Health (R01 MH077139 and RC2 MH089905), and the Sylvan C. Herman Foundation. Michel G. Nivard is supported by Royal Netherlands Academy of Science Professor Award (PAH/6635) to Dorret I. Boomsma. Veikko Salomaa was supported by the Finnish Foundation for Cardiovascular Research.

### COMPETING FINANCIAL INTERESTS

The authors declare no competing financial interests

## Online Methods

### Phenotype definition

We matched the original educational categories with the International Standard Classification of Education (ISCED), as described in **Supplementary Table 1**.Thereafter we used the equivalent of United States years of schooling to obtain the YOE. Going to college was defined as having an ISCED category>4.

To remove potential bias introduced by uncompleted education, we excluded all the individuals younger than 30 years at the time of sample collection.

### Sequencing procedures

Estonian WGS and Finnish WGS samples have been sequenced at Broad Institute on Illumina HiSeq X Ten machines run to 20x and 30x mean coverage (150bp paired reads), respectively. Estonian samples followed a PCR-free sample preparation. Swedish-WES and Finnish-WES samples were sequenced using either the Agilent SureSelect Human All Exon Kit or the Agilent SureSelect Human All Exon v.2 Kit. Sequencing was performed at Broad institute on Illumina GAII, Illumina HiSeq2000 or Illumina HiSeq X Ten. Mean target coverage was 90x.

All samples have been aligned against the GRCh37 human genome reference and BAM processing was carried out using BWA Picard. Genotype calling was done using GATK Haplotype Caller and was performed at Broad Institute for all studies.

### Hail software

To overcome the growing computational challenge of learning from large genomic datasets, we utilized Hail, an open-source software framework for scalably and flexibly analyzing such data (https://github.com/broadinstitute/hail). Hail, under active development, includes support for data import/export, quality control, analysis of population structure, and methods for performing both common and rare variant association. Hail is written in Scala (a Java virtual machine language) and builds on open-source software for scalable distributed computing including Hadoop (http://hadoop.apache.org/) and Spark (http://spark.apache.org/). Hail achieves nearperfect scalability for many tasks and can run on thousands of nodes. Hail automates fault-tolerant distribution of data and compute, greatly simplifying distributed pipeline execution compared to traditional HPC job schedulers like LSF and Grid Engine. Pipelines written in Hail’s high-level language typical require orders-of-magnitude fewer lines of code than comparable pipelines written in general purpose languages.

### Samples and variants QC

Quality control was performed independently for each study using *Hail.* We excluded individuals with high proportion of chimeric reads (>5%), high contamination (>5%) or an excessive number of singletons variants not observed in ExAC (> 100 for WES and>20,000 for WGS). We included only unrelated individuals (IBD proportion<0.2) and those for whom the sex predicted from genetic data matched the self-reported gender. We kept only ‘PASS’ variants, as determined by The Genome Analysis Toolkit^29^ Variant Quality Score Recalibration (VQSR) filter, are set to missing variants with GQ<20 and allele balance>0.8 or<0.2. We further excluded variants with call rate<0.8. In WGS data, we excluded low complexity regions as defined by Li ^30^. In the burden test analysis we excluded variants with both Hardy-Weinberg equilibrium test P-value<1×10^−6^ and negative inbreeding coefficient (expected heterozygosity less than observed heterozygosity).

### Annotation and URVs scores definition

Annotation was performed using SnpEff 4.2 (build 2015-12-05) ^31^ using Ensemble gene models from database GRCh37.75. We further annotated variants with SnpSift 4.2 (build 2015-12-05) ^32^ using annotations from database dbNSFP 2.9 ^33^. In **Supplementary Table S3** we have provided a detailed description of the criteria used for selecting variants in each score. The set of HC genes was defined separately for disruptive and damaging variants. For disruptive and synonymous mutations we defined HC genes those having a probability of being loss_of_function intolerant *(pLI)*>0.9 (N genes=3,488). For missense damaging mutation we used a missense *z-score*>3.09 (N genes=1,614)^5^. Both measures have been previously described^5^ and available online at ftp://ftp.broadinstitute.org/pub/ExAC_release/release0.3/functional_gene_constraint. We used a version derived from The Exome Aggregation Consortium without cases of psychiatric disorders.

### Principal component analysis and mixed models

We used a subset of high confidence SNPs to calculate principal components. We selected variants with minor allele frequency larger than 5%, call rate>90%, Hardy-Weinberg equilibrium test P-value> 1×10^−6^ and we pruned for variants in linkage disequilibrium using *plink* with command line ‘‐‐indep 50 5 2’.

We used a similar approach to filter variants used to generate the genetic relationship matrix (GRM). We then fit a liner mixed model including the GRM as random effect and age, sex, year of birth, (year of birth – 1950)^2^, (year of birth – 1950)^3^, the number of singletons synonymous variants not in ExAC and the number of URVs in HC genes as fixed effects.

### Association between URVs and educational attainment

We fit a linear regression model where the dependent variable was YOE and the independent predictors were: age, sex, year of birth, (year of birth – 1950)^2^, (year of birth – 1950)^3^, the 10 first principal components, the number of singletons synonymous variants not in ExAC, schizophrenia status (only in studies including schizophrenic patients) and the URV score (count of disruptive, damaging or synonymous URVs). We adjust for the number of all singleton synonymous variants to correct for potential technical artifacts.

### Brain-enrichment and brain-expression analysis

Using the Genotype-Tissue Expression consortia (GTEx) data ^23^, we ranked gene-expression levels (in RPKM) in brain tissues and defined the top 20% HC genes as “brain-expressed” (N. genes=683 and 313 for disruptive and damaging, respectively). Conversely, we defined “non brain expressed” the bottom 20% of the HC genes (N. genes=683 and 313 for disruptive and damaging, respectively).

We also compute estimated fold-change in the brain as follows. Suppose samples *1, 2,…, N_b_* are brain samples and samples *(N_b_+1), (N_b_+2),…, N*are the samples from other tissues. Denote with xij the expression of gene *j* and sample *i*, in reads per kilobase of transcript per million (RPKM). We compute fold-change (FC):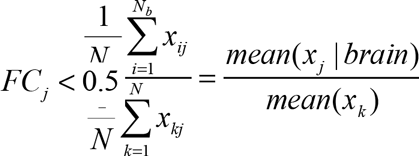 brain-enriched genes” and as “non-brain-enriched HC genes was 447 and 287 for disruptive and The number of non brain-enriched HC genes was 2,225 and 935 for disruptive and damaging mutations, respectively.

### Polygenic score, CNVs and runs homozygosity

The polygenic score for YOE was obtained from array data in the Swedish WES study (quality control for the array data have been previously described ^34^) and directly from WGS data in the Finnish-WGS and Estonian-WGS studies. We included all the independent markers with P-value<1 in largest GWAS of educational attainment ^16^ and obtained the polygenic score as weighted sum of risk alleles using the *score* command in Plink ^35^.

CNVs for the Swedish WES study were called as part of a separate project ^36^ using a composite pipeline comprising the CNV callers PennCNV, iPattern, Birdsuite and C-Score organized into component pipelines. We considered only rare CNVs by filtering out all CNVs that present at ≥ 1% allele frequency. CNVs<20kb or having fewer than 10 probes were also excluded. We used the plink *--cnv-intersect* function with a value of 0. 5 to determine the overlap between detected CNVs and the ldist of pathogenic CNVs reported in **Supplementary Table 4**.

CNVs in Finnish WGS and Estonian WGS were genotyped according to the methods described in ^37^ and implemented in Genome STRiP 2.0. Briefly, read depth information was collected from WGS data, excluding regions of the genome that are not uniquely alignable or have low sequence complexity, and adjusted for GC content bias. Each CNV reported in **Supplementary Table 4** was directly genotyped using Genome STRiP’s genotyping module, which examines the read depth across all samples and fits a constrained Gaussian mixture model with components representing each possible diploid copy number and sample-specific variance terms to account for differences in sequencing depth.

The summed runs of homozygosity were determined using the same pipeline described in ^7^. Specifically we used *plink* with command line ‘--homozyg --homozyg-window-snp 35 --homozyg-snp 35 --homozyg-kb 1500 --homozyg-gap 1000 --homozyg-density 250 -- homozyg-window-missing 5 --homozyg-window-het 1’.

### Gene-based burden test

We first extracted from each dataset variants falling within UCSC known genes and merged the four datasets using *plink*. If a variant was not present in all cohorts, we forced it as homozygous reference across the remaining cohorts (using “--fill-missing-a2” option in *plink*). We then computed principal components for the combined dataset after further merging with 1000 Genomes project samples as described in (Genovese et al, *jointly submitted).* To test the hypotheses that disruptive URVs in individual genes were associated with YOE and college status, we performed a burden test using the SKAT software ^38^ using default parameters (method=davies, impute.method=bestguess, r.corr=1.0), adjusting for age, sex, year of birth, (year of birth – 1950)^2^, (year of birth – 1950)^3^, the first 10 principal components, schizophrenia status and number of URVs identified in coding regions. We used a python wrapper to run the SKAT software (available at https://github.com/freeseek/gwaspipeline).

